# Charged sequence motifs increase affinity towards liquid-liquid phase separation

**DOI:** 10.1101/2021.09.08.459482

**Authors:** András László Szabó, Anna Sánta, Rita Pancsa, Zoltán Gáspári

## Abstract

Protein phase separation is a major governing factor in multiple cellular processes, such as those concerning RNA metabolism and RNA-binding proteins. Despite many key observations, the exact structural characteristics of proteins involved in the process are still not fully deciphered. In this work we show that proteins harbouring sequence regions with specific charged residue patterns are significantly associated with liquid-liquid phase separation. In particular, regions with repetitive arrays of alternating charges show the strongest association, whereas segments with generally high charge density and single α-helices also show detectable but weaker connections.

## Introduction

### Protein phase separation

Cells are continuously conducting various complex biochemical processes. These need to be efficiently performed and finely regulated in space and time, therefore cells employ a molecular process wherein macromolecules, mainly multi-domain proteins and RNAs, establish multivalent weak interactions to form functionally specialized liquid compartments, the so-called membraneless organelles (MLOs). Interestingly, this smart solution for the reversible and finely tuned compartmentalization of biochemical processes has only come to the focus of intensive research relatively recently. The general name for the phenomenon is phase separation, and it has been shown to have a role in critical processes of the living cell, such as chromatin regulation, RNA transcription, organization of the postsynaptic density etc. [1–3] Phase separation can result in different states of membraneless organelles (MLOs) from liquids through gels to solids. [4]

The molecular assemblies formed by these processes are typically micron-sized objects, containing multivalent proteins with multiple modular domains and disordered regions, often of low sequence complexity. [5] Regarding their functions, there are two main types of molecules participating in phase separation, as distinguished by Banani et al. [6], the “scaffolds” and the “clients”. The former type consists of molecules essential to the integrity of the MLOs. The latter type contains the majority of the components; however, they only participate in functionalities under certain circumstances. P bodies are a good example for this duality, they are scaffolded by a few critical RNA-binding proteins and store mRNAs and most protein components of the mRNA degradation machinery as clients. Scaffold-scaffold interactions are more persistent than scaffold-client interactions, and the composition changes according to a set of factors such as stress and the cell cycle. [1, 7]

The characteristic feature of liquid-liquid phase separation (LLPS) that distinguishes it from other processes related to phase separation is that the solution transitions into distinct phases where certain solutes are present in highly elevated concentrations, and these phases exhibit liquid-like properties. In this specific type of phenomenon, the term “scaffolds” is often replaced by “drivers”, referring to sets of proteins that are able to drive LLPS on their own. Smaller molecules and ions are omitted from this category, even if they are required for the initiation of the condensation process. Clients in this context are molecules that may partition into MLOs, but they hold no influence over their formation. [8]

A category of proteins whose members are especially prone to such processes are intrinsically disordered proteins (IDPs). Their low-complexity, prion-like sub-sequences govern LLPS, making the process prone to undergo material state transitions, such as the liquid-solid transition exhibited by RNA-binding protein FUS as well as TAR DNA-binding protein 43 (TDP-43). Liquid-solid phase transitions – in specific cases called aggregation – are often associated with severe diseases such as amyotrophic lateral sclerosis (ALS), making the examination of IDPs and LLPS rigorously researched fields. [9, 10]

Pak et al. [11] described the phase separation of the intrinsically disordered intracellular domain of Nephrin (NICD) as complex coacervation. When expressed as a soluble protein, NICD formed droplets in HeLa cells, and LLPS was also observed in vitro when mixed with positively supercharged GFP. While no specific sequence was identified in NICD responsible for LLPS, a pattern of blocks of negatively charged residues was observed. This implies phase separation to be robust in withstanding mutations, as shuffle and deletion mutants were still able to drive phase separation as long as charged blocks were retained.

Many proteins involved in the phase separation phenomena have been identified in the last 20 years, and thus it has become a relevant issue to catalogue them. PhaSepDB [12] is a manually curated database that has been created with the specific purpose of providing researchers with a detailed, reliable collection of information about the proteins connected to the phenomenon. There are other, more specific databases of phase-separating protein such as the stress granule protein database constructed by Nunes et al. [13], or PhaSePro that specifically collects LLPS drivers [14], but as of today one of the most comprehensive databases is PhaSepDB. Data collection and processing for such a library of proteins consists of several levels, and new entries are added into the database after their respective publications show evidence on the localization of the proteins to MLOs.

There are three categories of protein sets available in PhaSepDB: The first one is the “Reviewed” category that only contains articles published after January 1st of 2000, constituting the most reliable subset. The second category is called “UniProt reviewed” that includes a wider range of articles, not restricted by publication date, although its results must be confirmed by more recent studies. The third and final category is the “High-throughput” research data, generated by high-throughput methods categorized as either organelle purification, proximity labelling, immunofluorescence image-based screen or affinity purification. We have used the Reviewed database that was released in 2019, consisting of 234 human entries.

### Single α-Helices and other charged residue repeats

Single α-helices (SAHs) are protein segments that form stable and rigid helical structures even in isolation. [15, 16] These regions are rich in arginine, lysine and glutamate and exhibit a characteristic repetitive pattern of oppositely charged residues such as glutamic acid and lysine. Their length may vary from a few dozen up to about 200 residues, and they are stabilized by intrahelical salt bridges. [17] These regions show similar characteristics to coiled coils, as Peckham et al. [18] showed that ~4% of human proteins that were previously predicted to contain coiled coils actually have SAHs instead. SAH domains have been found to be prevalent in RNA-binding proteins [15], although the SAH region itself is unlikely to directly contribute to the interaction with RNA.

Inspired by the prominent role of RNAs and RNA-binding proteins in phase separation, we intended to investigate whether proteins involved in LLPS are enriched in SAH segments. As SAH segments constitute a special case of sequences with high charge density, we have extended our analysis to segments with repeating patterns of charged residues, termed charged residue repeats, CRRs, as well as with segments enriched in charged residues but without any further specifications, denoted charge-dense regions, CDRs, below. We differentiate between “unsigned” and “signed” CRRs on the basis of their net charge. In this classification, most CRRs are special cases of CDRs, and SAHs constitute a sub-class of CRRs (Figure 1).

**Figure 1.**
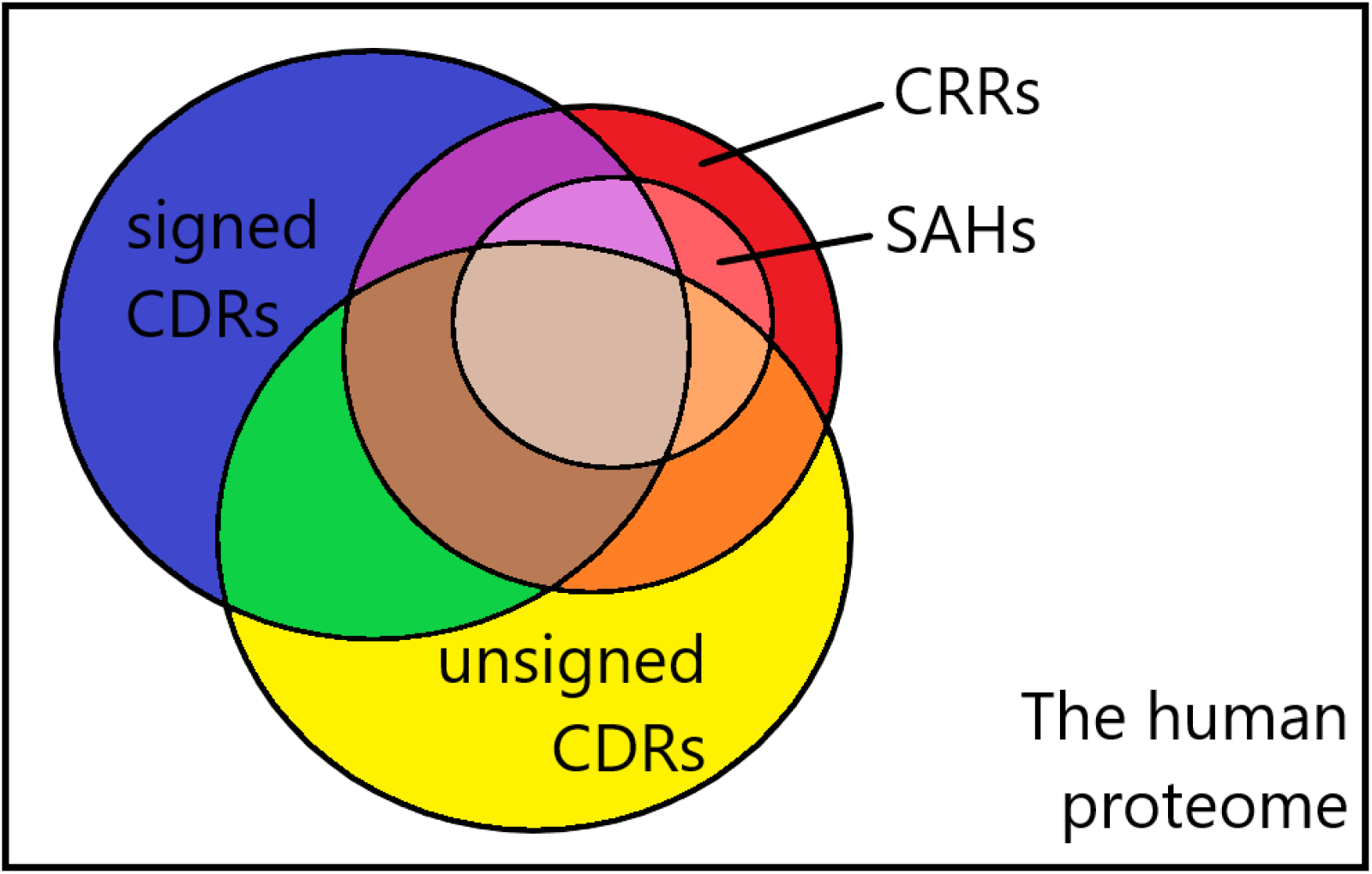
Venn diagram illustrating the relations between single α-helices, charged residue repeats, as well as signed and unsigned charge-dense regions within the human proteome.

It is important to note that while the majority of CRRs satisfy at least one of the two definitions for charge-dense regions, not all of them fall under those categories. The cleavage stimulation factor subunit 2 (CSTF2, UniProtKB ID: P33240) for example contains 12 × 5 AA tandem repeats on a 60 residue-long segment that was detected as a CRR by the FT_CHARGE algorithm - a 105 residue-long sub-sequence encompassing the annotated region was highlighted (Figure 2).

**Figure 2.**
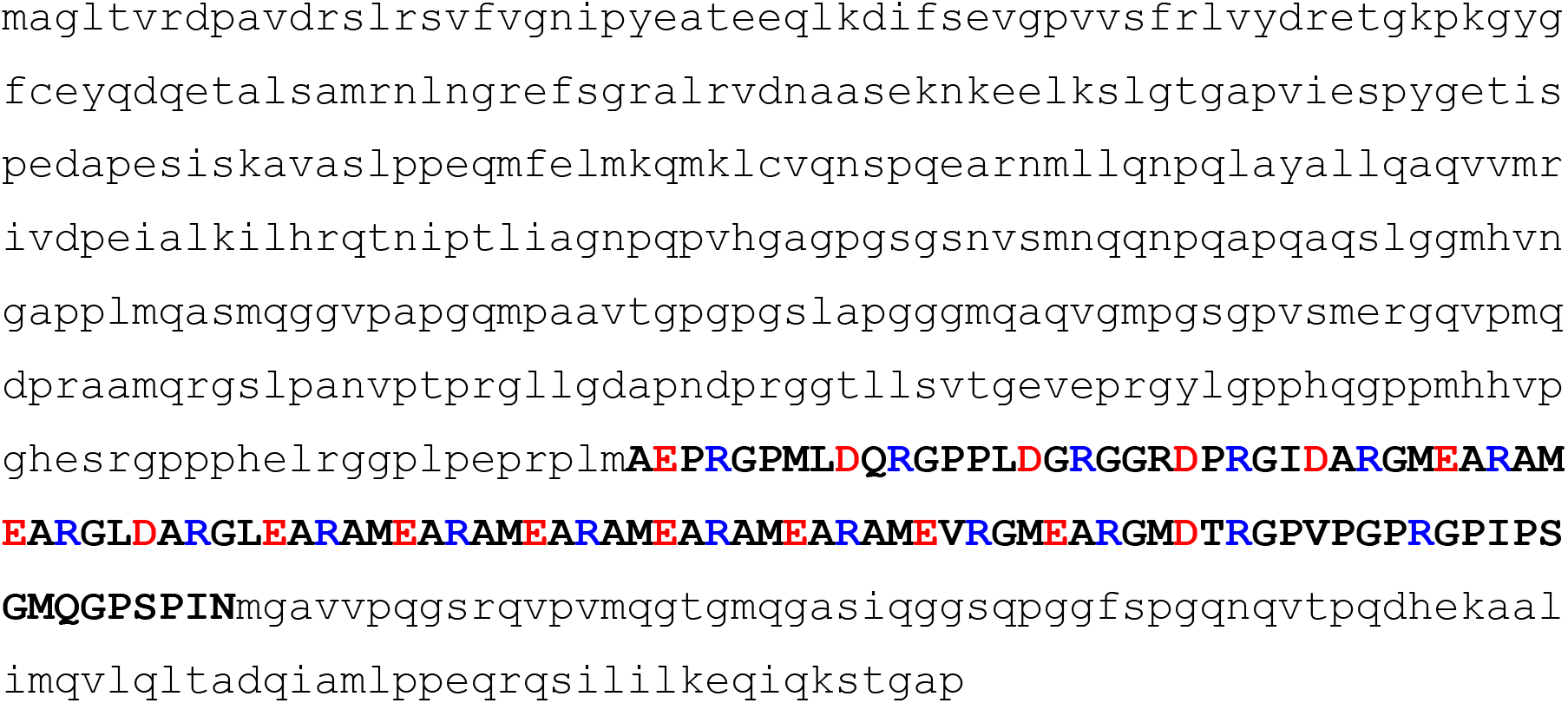
The charged repeat region of protein CSTF2 that is not recognized as a SAH.

The region does not qualify as a SAH due to its slightly higher Fourier frequency, although there are multiple helical structures along the entire sequence of this protein. While up to this point, we could not find a region that qualifies as a SAH while remaining outside our definitions for CDRs, we would reserve the possibility for their existence as the herein applied representation of the human proteome only included a single isoform per protein.

To strengthen the specificity of our investigations, we have performed all our evaluations by excluding proteins with transmembrane segments. The rationale behind this is that the presence of charged regions is expected to be characteristic for soluble proteins. Without excluding transmembrane proteins, any possible association between charged regions and LLPS might simply reflect that phase separation is primarily occurring between soluble biomacromolecules.

## Results

### CDRs, CRRs and SAHs in the human proteome

We have restricted our studies to the human proteome as specified at the beginning of the Methods section. For the prediction of SAHs, we used our previously established methodology, involving the FT_CHARGE algorithm. CRR detection was carried out by relaxing the requirement for the charge frequency characteristic for SAHs. CDRs were detected by an algorithm enumerating Asp, Glu, Arg and Lys residues in sequence windows of different length (see Methods for details). Since the vast majority of human proteins was found to contain a CDR according to our original detection threshold, we have applied a stricter criterion and investigated only the top 1% of CDRs (see Supplementary Table S1).

### CRR-containing proteins show significant enrichment in proteins involved in LLPS, while proteins with SAHs show a weaker but still detectable association

Fisher’s exact test of independence showed that there is an exceptionally high probability that charged residue repeats (CRRs) contribute to the LLPS of human proteins (Figure 3, Supplementary Figure S1, as well as Tables S1 and S2).

**Figure 3.**
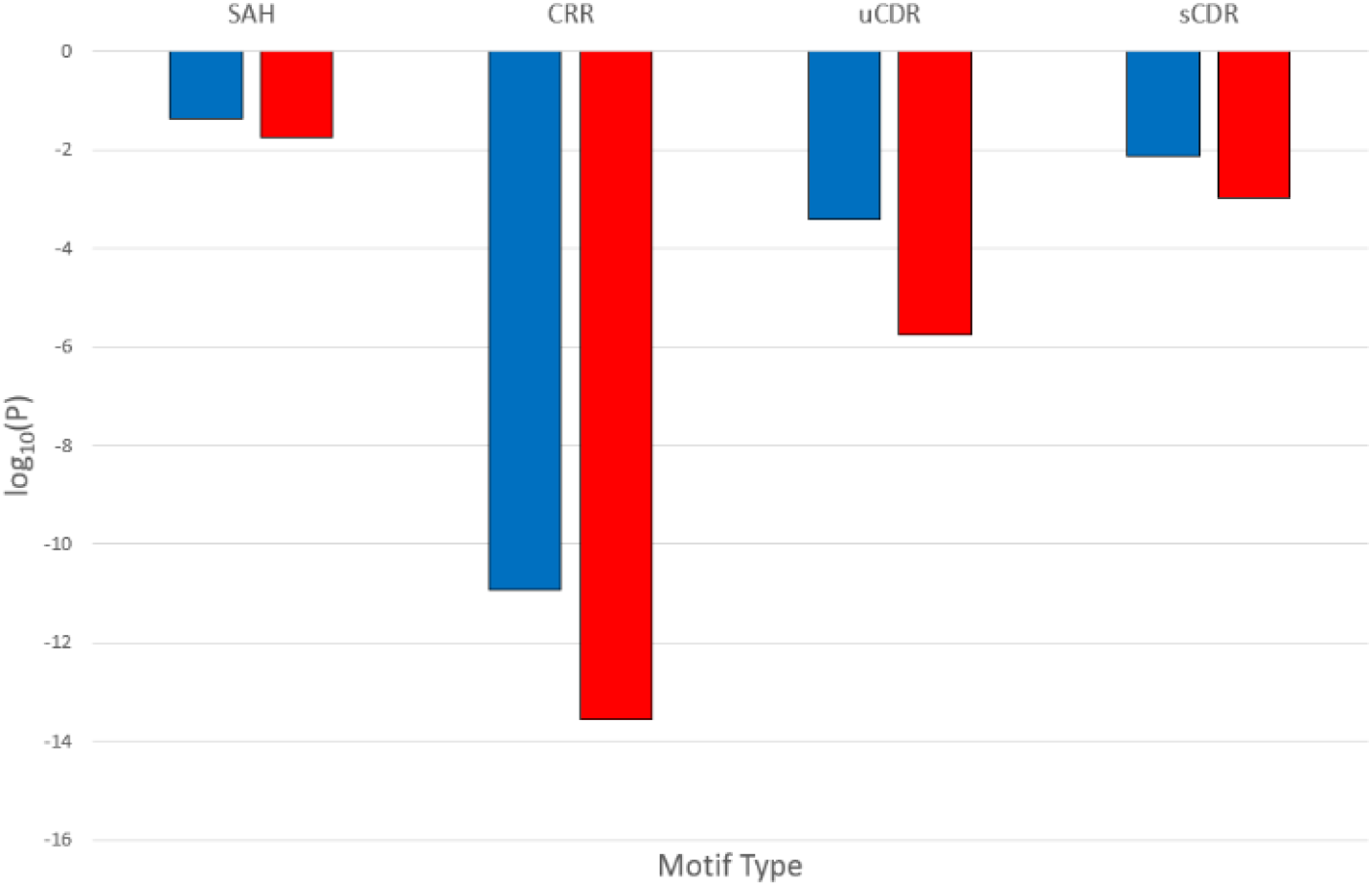
Decimal logarithm of the P-values that belong to Fisher’s tests where the functional attribute is the protein’s relation to LLPS, and the structural attribute is the presence of different sequence motifs. Tests involving the entire proteome are showcased in red while tests that excluded transmembrane proteins are in blue.

SAHs can be regarded as special subsets of CRRs detected with FT_CHARGE with the frequency region between 1/9 and 1/6. SAH regions still exhibit a statistically significant enrichment in LLPS-associated proteins, although this is considerably weaker than for CRRs in general (Figure 4 and Supplementary Tables S1 and S2). Another way to illustrate the level of association between LLPS and the presence of sequence motifs is to compare their enrichment in LLPS-related proteins. For example, the ratio of sequences with CRRs to other proteins is 1010:19418 in the full proteome in case of entries that are unrelated to LLPS. The same ratio in case of related sequences is 44:187. Which means that the ratio increases 4.52-fold between the two functional categories (Figure 5).

**Figure 4.**
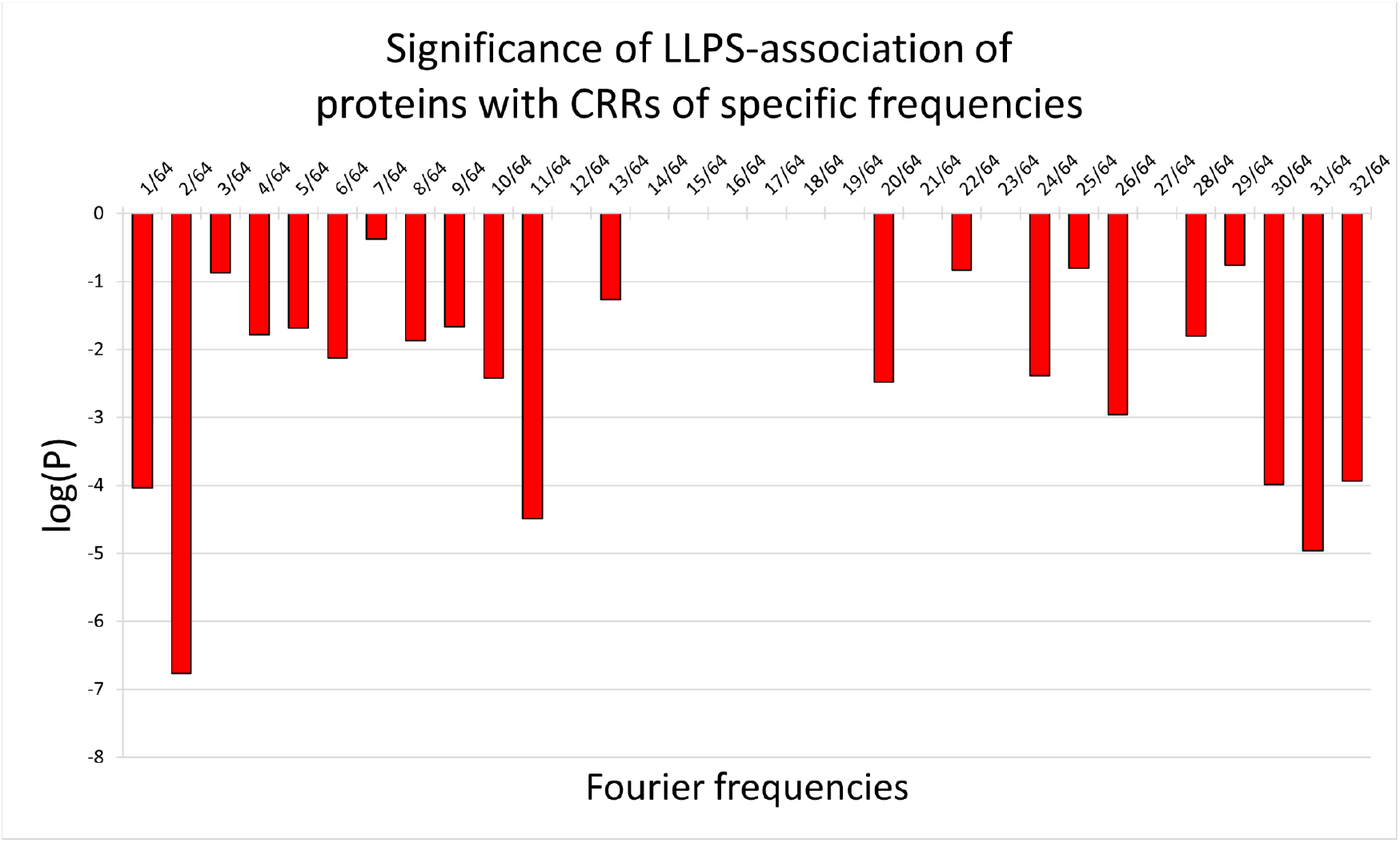
Decimal logarithm of the P-values that belong to Fisher’s tests where the functional attribute is the protein’s relation to LLPS, and the structural attribute is the presence of charged residue repeats with specific Fourier frequencies.

**Figure 5.**
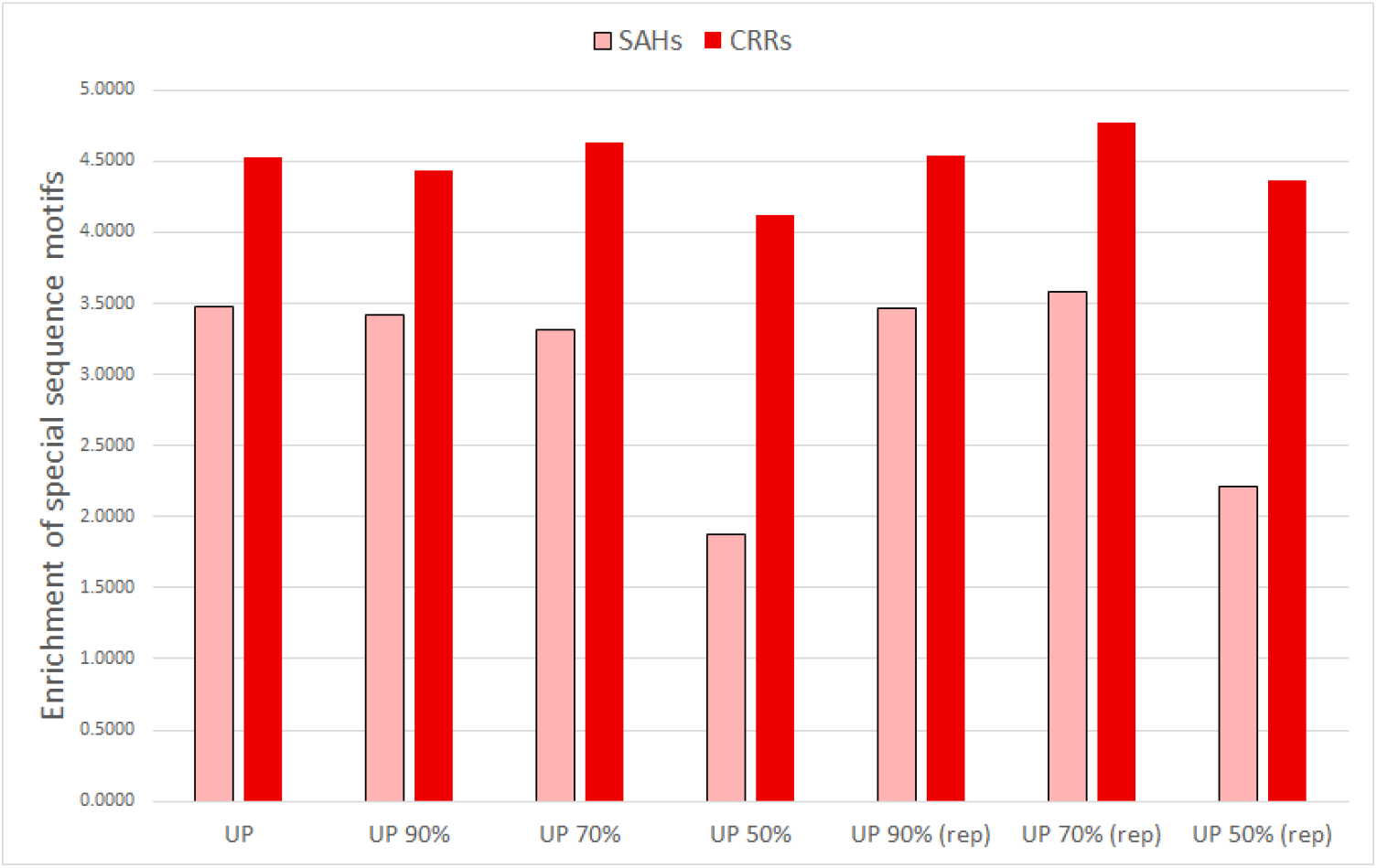
CRR-enrichment (proportional increase of the ratio of CRR-containing sequences) in case of the full proteome as well as its variants with reduced redundancy (with clusters and with representative sequences only, see Methods for details).

### Regions enriched in charged residues in general are prevalent in proteins associated with LLPS

We have investigated whether regions enriched in charged residues but not necessarily exhibiting regular repeating patterns are also associated with LLPS. To this end, besides the charged residue repeats detected by the FT_CHARGE algorithm, a more general type of protein sub-sequences called “Charge-Dense Regions / CDRs” was similarly probed for relations to LLPS. We have formulated two different approaches for their detection. One of them defines CDRs as protein sub-sequences where the density of charged residues is considerably higher than average. The other one defines them as protein sub-sequences with an overall charge the absolute value of which is significantly higher than average. We refer to the regions gained through the former approach as “unsigned CDRs”, while regions resulting from using the other definition were named “signed CDR” - as their charges may have a negative sign, too. The protocols for detecting such components are described in detail in the Method section.

### Unsigned and signed CDRs

If we consider the top ~1% of all hits to be charge-dense regions, then approximately 47.10% of human proteins contain some kind of charge-dense region. When these CDRs are compared to CRRs, it turns out that 90.99% of charged residue repeats display an above 90% overlap with CDRs, with 87.80% of all CRRs being entirely encompassed by charge-dense regions, while 2.53% of them are free from any overlap. Sequence-wise, 15.92% of the human proteome is categorized as CDRs with the above-mentioned parameters.

Considering the top 1% of all hits as signed CDRs results in 22.76% of charged residue repeats having at least 90% overlap with the top 1% of CDRs, and 20.25% having total coverage, while 37.14% having no coverage at all.

### ROC analysis of motif presence as predictor of LLPS

We have performed receiver operating characteristic (ROC) analysis to assess whether the presence of such motifs has predictive value for LLPS-association for a given protein. We found that while the presence of these motifs is not a strong predictor of LLPS, the absence of such regions largely precludes the participation of the protein to form condensates (Figure 6).

**Figure 6.**
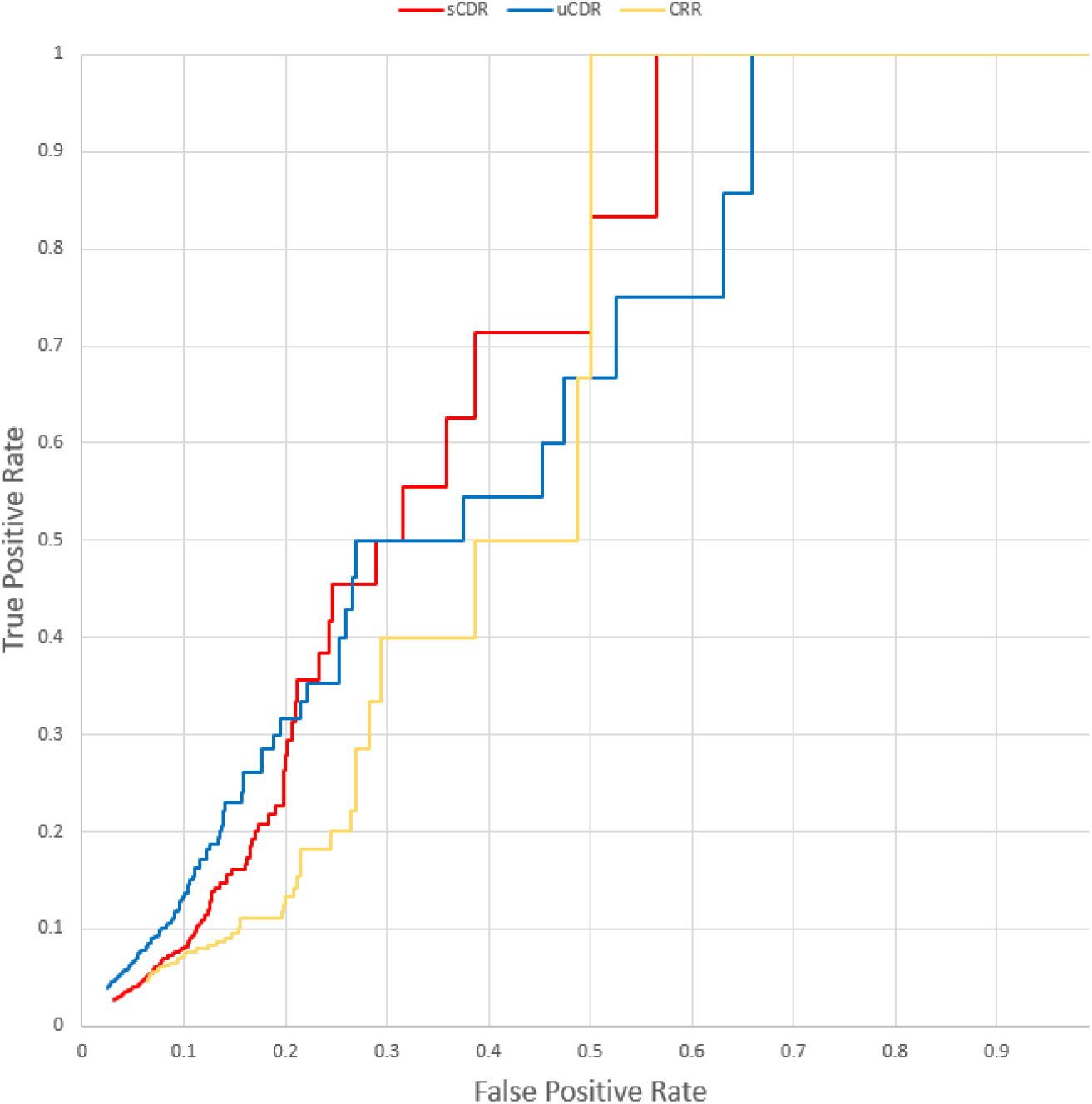
Receiver operating characteristic (ROC) tests on CRRs, unsigned CDRs and signed CDRs in non-transmembrane proteins, evaluating the presence of these motifs as an indicator of affinity towards LLPS. For CRRs the test shows the highest value at 0.5 FPR (TPR = 1) when the threshold is set to 0.38, meaning that sequences where at least 0.38 of all residues are assigned to charged residue repeats exhibit a high probability of participating in LLPS. These numbers are TPR = 0.667 and threshold = 0.80 for uCDRs, and TPR = 0.714 and threshold = 0.83 for sCDRs. The ROC curves suggest that the presence of CDRs/CRRs is not a strong predictor for LLPS, but the absence of such motifs is an indicator that the protein is not associated with LLPS.

### Robustness of the motif-LLPS association in random sequence sets

The robustness of the association was also investigated in random sequence sets generated to match the size distribution of proteins in PhaSepDB (see Methods). Our observations confirm that these motifs are specifically enriched in LLPS-associated proteins (Supplementary Figure S2 and Table S4).

### Functional analysis of charged motif-containing proteins in relation to LLPS

The Overrepresentation Test (OT) of Panther’s Gene List Analysis tools was also utilized to compare the various special motifs in sequences that are related to LLPS versus the regions of proteins that are unrelated to the phenomenon. A total of eight tests were carried out, each comparing either LLPS-related or LLPS-unrelated SAHs, CRRs and signed as well as unsigned CDRs. From the OT-given Gene Ontology (GO) terms we manually selected the ones that are at the lowest level within their respective branch of the GO term hierarchy, meaning that only the most specific - and thus least redundant - terms were analysed further. These terms were compiled into a table where each term’s enrichment - or lack thereof - was noted in case of the four different kinds of LLPS-related and LLPS-unrelated motifs. From there it is possible to categorize each term, based on the number of LLPS-related categories they were enriched in (0-4) and the number of LLPS-unrelated categories they were enriched in (0-4). This method of categorization created a gradient table that shows the distribution of GO terms from highly related to LLPS to highly unrelated to LLPS. The table then could be reduced to contain GO terms with specific keywords in them, such as ‘regulation’, ‘RNA’ or ‘signal’ (Table 2, Supplementary Table S3).

**Table 1.** Summary table exhibiting the amount of sequences associated with different sequence motifs, as well as the fraction of residues encompassed by those motifs. Includes data for the full proteome and its variants with reduced redundancy.

| Number of proteins |  |  |  |  |
| --- | --- | --- | --- | --- |
| Sequence motifs | Full proteome | Redundancy-filtered proteome |  |  |
|  |  | 90% | 70% | 50% |
| All proteins | 20 659 | 19 638 | 18 294 | 15 672 |
| CDRs (unsigned) | 9 731 | 9 314 | 8 792 | 7 669 |
| CDRs (signed) | 14 065 | 13 471 | 12 757 | 11 097 |
| CRRs (all) | 1 054 | 1 025 | 985 | 910 |
| CRRs ($\geq 90\%$ overlap with uCDRs) | 782 | 763 | 738 | 688 |
| CRRs ($\geq 90\%$ overlap with sCDRs) | 227 | 221 | 214 | 208 |
| SAHs (all) | 134 | 131 | 126 | 118 |
| Percentage of residues |  |  |  |  |
| All proteins | 100% | 96.74% | 92.04% | 80.73% |
| CDRs (unsigned) | 10.02% | 9.63% | 9.16% | 8.22% |
| CDRs (signed) | 14.53% | 13.98% | 13.30% | 11.58% |
| CRRs (all) | 0.82% | 0.80% | 0.77% | 0.72% |
| CRRs ($\geq 90\%$ overlap with uCDRs) | 0.54% | 0.52% | 0.50% | 0.47% |
| CRRs ($\geq 90\%$ overlap with sCDRs) | 0.14% | 0.13% | 0.13% | 0.12% |
| SAHs (all) | 0.11% | 0.11% | 0.10% | 0.10% |

**Table 2.** Gradient tables depicting the distribution of all GO terms (a), terms related to RNAs (b), terms related to responses (c), terms related to metabolism (d). In the latter case the actual keyword was ‘metabol’ to include GO terms containing ‘metabolism’ and ‘metabolic’ as well, much like the keyword ‘RNA’ encompasses ‘mRNA’, ‘rRNA’, ‘snoRNA’, ‘RNA-binding’, etc.

|  |  |  |  |  |  |
| --- | --- | --- | --- | --- | --- |
| a) | Total number of entries = |  |  |  | 307 |
| Enrichment | 0/4 Unrelated | 1/4 Unrelated | 2/4 Unrelated | 3/4 Unrelated | 4/4 Unrelated |
| 4/4 Related | 1 | 0 | 0 | 0 | 0 |
| 3/4 Related | 10 | 0 | 0 | 0 | 0 |
| 2/4 Related | 53 | 7 | 1 | 0 | 0 |
| 1/4 Related | 90 | 5 | 3 | 0 | 0 |
| 0/4 Related | 9 | 112 | 16 | 0 | 0 |
| b) | Total number of entries related to 'RNA' = |  |  |  | 55 |
| Enrichment | 0/4 Unrelated | 1/4 Unrelated | 2/4 Unrelated | 3/4 Unrelated | 4/4 Unrelated |
| 4/4 Related | 0 | 0 | 0 | 0 | 0 |
| 3/4 Related | 5 | 0 | 0 | 0 | 0 |
| 2/4 Related | 13 | 1 | 0 | 0 | 0 |
| 1/4 Related | 26 | 0 | 3 | 0 | 0 |
| 0/4 Related | 0 | 7 | 0 | 0 | 0 |
| c) | Total number of entries related to 'response' = |  |  |  | 28 |
| Enrichment | 0/4 Unrelated | 1/4 Unrelated | 2/4 Unrelated | 3/4 Unrelated | 4/4 Unrelated |
| 4/4 Related | 0 | 0 | 0 | 0 | 0 |
| 3/4 Related | 0 | 0 | 0 | 0 | 0 |
| 2/4 Related | 10 | 0 | 0 | 0 | 0 |
| 1/4 Related | 9 | 0 | 0 | 0 | 0 |
| 0/4 Related | 1 | 6 | 2 | 0 | 0 |
| d) | Total number of entries related to 'metabol' = |  |  |  | 22 |
| Enrichment | 0/4 Unrelated | 1/4 Unrelated | 2/4 Unrelated | 3/4 Unrelated | 4/4 Unrelated |
| 4/4 Related | 0 | 0 | 0 | 0 | 0 |
| 3/4 Related | 0 | 0 | 0 | 0 | 0 |
| 2/4 Related | 1 | 0 | 0 | 0 | 0 |
| 1/4 Related | 2 | 1 | 0 | 0 | 0 |
| 0/4 Related | 4 | 14 | 0 | 0 | 0 |

A simple analysis of the relationship between postsynaptic localization, RNA-binding function, presence of charged motifs and LLPS-association (Figure 7 and Supplementary Figures S3-S9) reveal that there are barely any proteins related to either LLPS, the postsynaptic density or the molecular function of RNA-binding that are devoid of any motifs with either charged patterns or high charge density. Furthermore, the distributions implicate that RNA-binding proteins are more likely to participate in phase separation than PSD components as the ratio of entries in the middle (yellow) section doubles between the two figures. This enrichment can also be observed between PSD proteins and RNA-binding sequences when we only consider CRRs as a structural factor. Although the number of cases is small, it also seems that for RNA-binding proteins, the presence of charged motifs / CRRs is more prevalent when considering proteins participating in LLPS.

**Figure 7.**
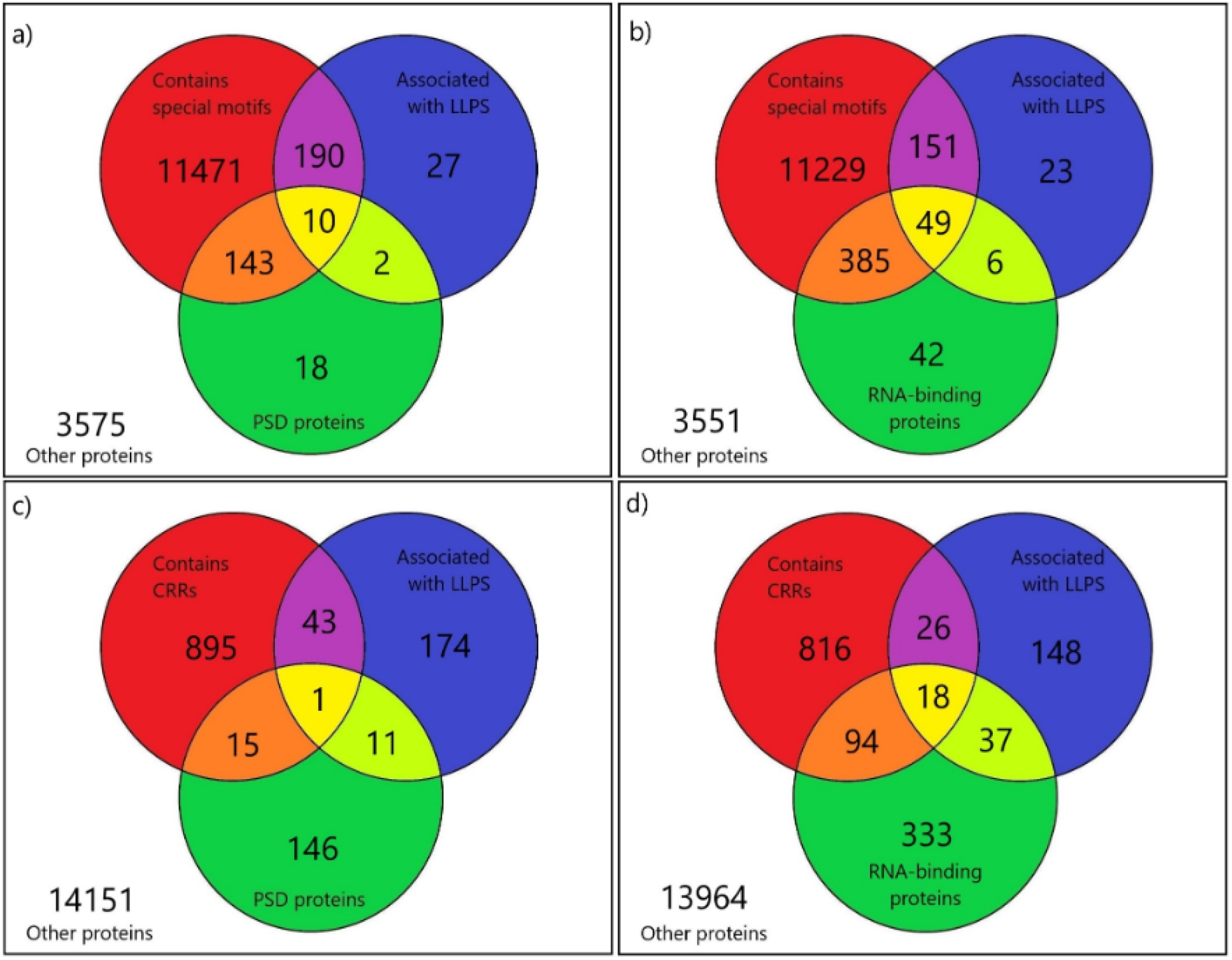
Distribution of human, non-transmembrane proteins according to their special charged motif (a-b) or CRR (c-d) content, LLPS association and their functional role in the PSD (a, c) or RNA binding (b, d).

## Discussion

### Case studies

Our results clearly indicate that proteins with regions containing a high fraction of charged residues are preferentially associated with LLPS. Our approach allowed us to investigate multiple aspects of this association, consideration of the presence or absence of specific repetitive patterns. Below we discuss several proteins, the possible relationship between LLPS and the presence of SAHs/CRRs/CDRs. Examples, wherever it is possible, are based on the information available in the richly annotated PhaSePro database, where information about the region responsible for LLPS is provided if available.

Table 1. contained 1054 protein sequences that encompass CRRs but out of those only 44 sequences are considered to be LLPS-related. These sequences were investigated further to explore their connection to phase separation. The investigation revealed that a large variety of proteins may participate in the phenomenon, including ribonucleases [19], splicing factors [20, 21], transcriptional repressors [22, 23], translational initiation factors [24], transport-[25, 26] and Zinc finger proteins [27, 28], etc. However, there happens to be a common feature in most of the investigated proteins, as 31 of them carry out the molecular function of RNA-binding. [19–37] The fact that 70.45% of these proteins can bind RNA molecules is significant and reinforces that RNA-binding is a process that is promoted by charged protein regions, such as CRRs. Another plausible explanation may be that many membrane-less organelles – that are formed by LLPS – contain RNAs. According to the frequencies of the CRRs predicted by FT_CHARGE, 5/44 sequences possibly contain single α-helices, all of which constitute at least 10% of the respective sequence.

It is important to note that while RNA-binding connects most of these proteins, their sequences as well as their biological function vary greatly. There were only two cases where multiple sequences got clustered together. In the first case the two proteins were the probable global transcription activator SNF2L2 and the transcription activator BRG1. Both are involved in transcriptional activation and repression of select genes by chromatin remodelling. [38] They are also components of SWI/SNF chromatin remodelling complexes that carry out key enzymatic activities and change chromatin structure. While these two proteins proved to be participating in the same processes, they still only showed 59.75% sequence identity.

In the second case there were three proteins in the same cluster, proline- and glutamine-rich splicing factor SFPQ being the representative sequence, non-POU domain-containing octamer-binding protein NONO showing 58.39% identity, and paraspeckle component 1 PSPC1 exhibiting 54.49% sequence identity. The three proteins cooperatively regulate androgen receptor-mediated gene transcription activity in the Sertoli cell line [39, 40] and, besides the noncoding RNA NEAT1, are major components of the paraspeckle. These proteins contain a conserved SAH located at the C-terminal end of a right-handed coiled coil segment. [41] The region responsible for dimerization, and thus LLPS, is the so-called NOPS region on the N-terminal side of the coiled coil. The same goes for the trinucleotide repeat-containing gene 6B protein (TNRC6B) that participates in RNA-mediated gene silencing and acts as a scaffolding protein that can simultaneously interact with argonaute proteins regulating miRNA and siRNA maturation and recruit deadenylase complexes carrying out poly(A) tail-shortening processes. [42] Finally, the E3 ubiquitin-protein ligase RNF168 accumulates repair proteins to sites of DNA-damage via binding to ubiquitinated histone H2A and H2AX. [43] Apart from SNF2L2 and BRG1, all the above proteins do encompass at least one alpha helical structure, and all of them seem to exhibit some kind of RNA-binding trait, except RNF168.

SynGAP1 is part of the NMDAR complexes in excitatory synapses. The mouse ortholog has been explicitly shown to participate in LLPS, with the responsible region being its C-terminal segment forming a trimeric coiled coil and having a PDZ-domain binding motif that interacts with PSD-95. The role of SynGAP in LLPS formation in the postsynaptic density has been demonstrated in multiple experiments. The key feature for LLPS in SynGAP is its multivalent nature provided by the trimerization via its coiled coil motif. SynGAP does not contain any SAHs or CRRs detected by our methods, but it bears CDRs and so does its partner protein PSD-95 — a.k.a. Disks large homolog 4. This interaction is part of a larger scaffolding protein network in the PSD involving DLGAP1 (the guanylate kinase-associated protein GKAP — a.k.a. Disks large-associated protein 1), Shank3 (SH3 and multiple ankyrin repeat domains proteins), Homer3 and the C-terminal region of the NR2B subunit of NMDAR. Mixed all together, they demonstrated phase separation *in vitro*, driven by a complex set of multivalent interactions. [44] All of the aforementioned PSD proteins contain CDRs, but neither SAHs or other CRRs, and only SynGAP and PSD-95 are found in PhaSePro, where in both cases, the LLPS-driving region is overlapping with at least one CDR (see Figure 8). It should be noted, that unlike in the previously mentioned networks, none of these proteins are associated with nucleic acid binding. LLPS occurs by the cell membrane, forming a PSD condensate attached to NMDAR clusters. This group is rich in protein-protein interaction domains, such as SH3, PDZ, SAM domains and their binding motifs.

**Figure 8.**
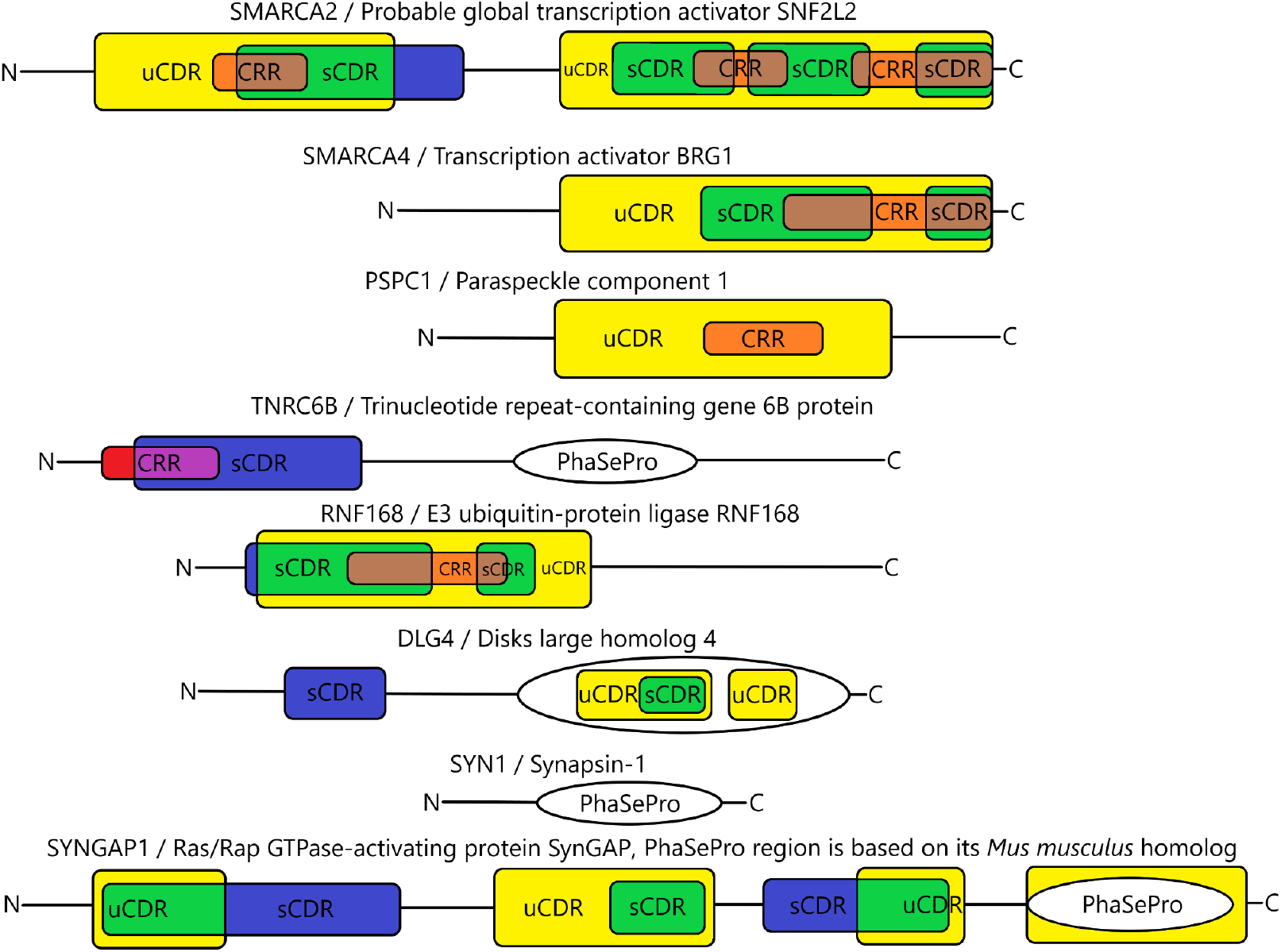
Sequence motifs as well as PhaSePro-based sub-sequences that drive LLPS. The colour-coding of individual motifs as well as their overlaps are in accordance with Figure 1. While currently there isn’t any experimental evidence that the human protein SynGAP participates in the phenomenon, its *Mus musculus* homolog has, thus a PhaSePro region has been annotated according to that.

Synapsin-1 participates in neurotransmitter release in the presynaptic region. The region responsible for phase separation is its Pro/Gln-rich C-terminal half and the protein can form condensates on its own or with binding partners containing SH3 domains. The condensates can also contain synaptic vesicles (SVs), providing the basis of self-organization of SV clusters. Phosphorylation of Synapsin-1 by CamKII causes dissociation of the condensates. [45] Synapsin-1 does not seem to contain any SAHs/CRRs/CDRs.

In the case of the 44 sequences that are related to LLPS and contain CRRs, it has also been observed that only one of them (U3 small nucleolar RNA-associated protein 6 homolog, UniProtKB ID: Q9NYH9) is devoid of predicted disordered segments according to their UniProtKB annotation. Furthermore, from the remaining 43 sequences there are only two (Paraspeckle component 1, UniProtKB ID: Q8WXF1 and Non-POU domain-containing octamer-binding protein, UniProtKB ID: Q15233) where none of the disordered segments overlap with any of the charged regions (Figure 8 and Supplementary Table S5). It must be noted here that there are disordered segments within the other 41 sequences that do not overlap with any of the charged regions but in each of those 41 sequences there is at least one disordered segment that does.

While these observations may indicate that disordered regions played a role in at least some of the cases where CRRs were associated with LLPS it must be urged that there are several proteins with experimental evidence where phase separation is directly driven by electrostatic interactions. There are fourteen such sequences within the PhaSePro database, out of which nine are human entries. Either CRRs, uCDRs and sCDRs are assigned to all but one of these human entries, the probable ATP-dependent RNA helicase DDX4 (UniProtKB ID: Q9NQI0) that contains at least one region with a charged density just below the threshold to be included in the top 1% of CDRs.

### Role of charged regions in LLPS

While our results clearly demonstrate an association between charged segments and LLPS, the rationale behind this is not straightforward. The association between LLPS and charged motifs weakens but still remains significant when we exclude transmembrane proteins. It should be noted that while this decision is based on the general expectation that primarily soluble proteins contain charged motifs and participate in phase separation, there are examples of transmembrane proteins involved in LLPS. In our reference proteome two transmembrane proteins are associated with LLPS: Linker for activation of T-cells family member 1 / LAT (UniProtKB ID: O43561) and Nephrin (UniProtKB ID: O60500). We have also identified several transmembrane proteins, both single- and multi-pass ones (e.g., Vesicle-associated membrane protein-associated proteins, sodium bicarbonate transporter 3 etc.), that contain at least one SAH, the most specific charged motif investigated here. Thus, the inclusion of transmembrane proteins is also not necessarily a biologically unfeasible approach.

We have also investigated whether proteins of similar length to those in PhaSepDB exhibit a distribution of charged elements characteristic of proteins involved in LLPS. Our results suggest that all of the investigated motifs, but especially CRRs and uCDRs are more abundant in LLPS-associated proteins than in our random datasets. These results suggest that the observed association between charged motifs and LLPS is unlikely to reflect some trivial biophysical constraint (see Supplementary Table S7 and Figure S9).

Importantly, our detailed investigation of the association between LLPS and charged motifs reveals that the absence of such motifs largely precludes the participation of a protein in LLPS and not that the presence of a CDR, CRR or SAH would be a strong indicator of LLPS formation. Here it should be noted that the accuracy of listings of proteins associated with LLPS is expected to increase with novel proteins recognized and curation efforts getting more intensive, probably resulting in an overall increase of proteins reliably proven to form condensates. That said, it is highly unlikely that charged regions identified here in this study might be directly responsible for LLPS in general. It is not the case for all proteins detailed above, let alone phase separating proteins without a CRR or CDR. Thus, it is more likely that the enrichment of CRRs and CDRs in LLPS-prone proteins has a more complex explanation. It should be noted that little is known about the structural features of CRRs and CDRs. Not surprisingly, many regions rich in charged residues, including SAHs, are predicted to be intrinsically disordered by several prediction methods. However, SAHs are known to possess a stable well-characterized structure and can be recognized by specialized algorithms. Thus, the presence of specific structural preferences for other types of CRRs cannot be completely excluded, and the formation of more or less well-defined monomeric or oligomeric structures might occur for specific sequences. Regularly alternating blocks of positively and negatively charged residues might offer a platform for intermolecular associations in multiple mutual orientations. The mediator of RNA polymerase II transcription subunit 1 (MED1, UniProtKB ID: Q15648) is a great example for this, as it has a relatively large region (Figure 9) experimentally connected to phase separation that contains acidic and basic segments. [46] This intrinsically disordered region also overlaps with CRR, uCDR and sCDR predictions.

**Figure 9.**
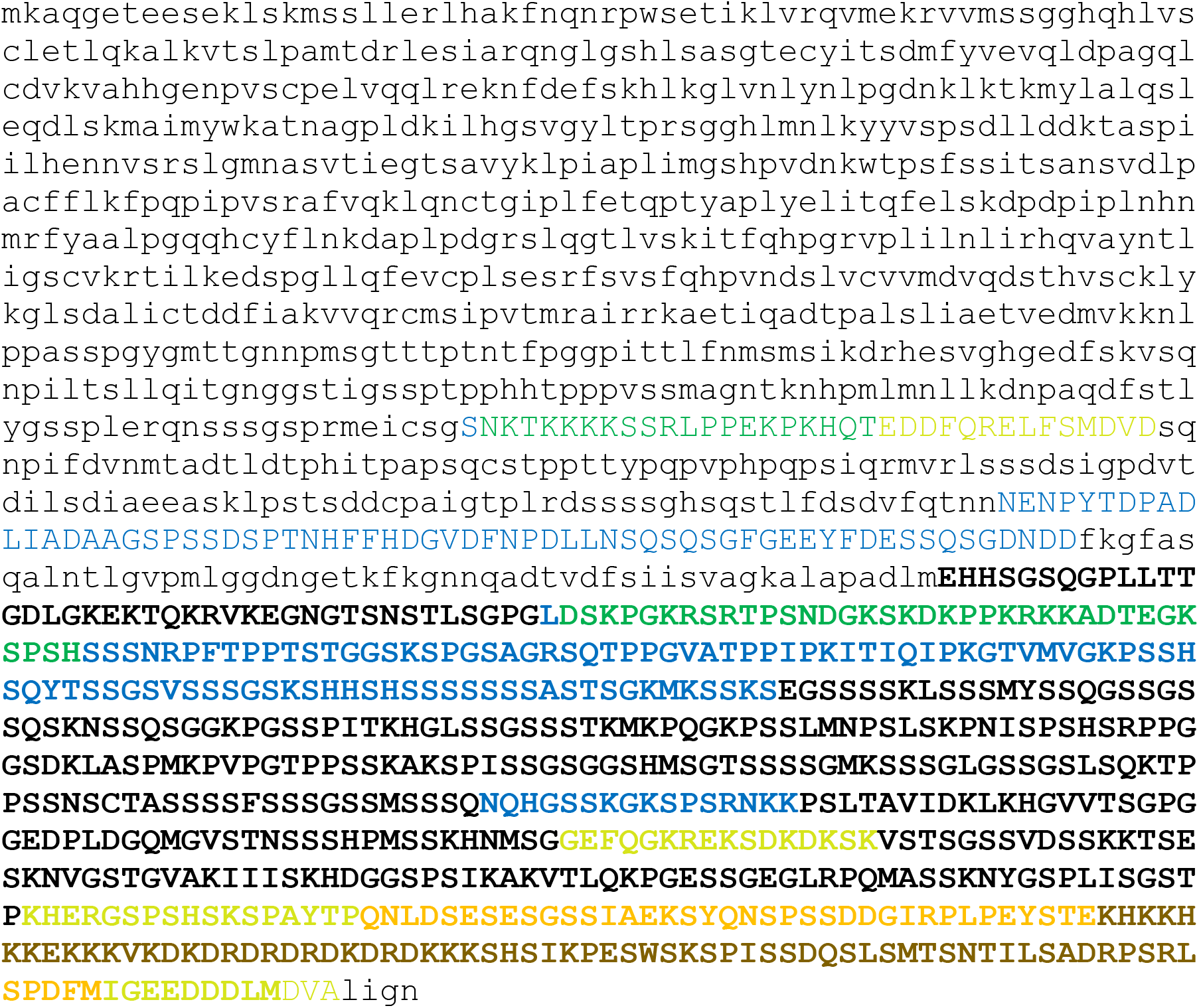
Sequence of MED1 colored according to the presence of charged motifs (color-coded according to Figure 1). The region involved in phase separation is highlighted in bold.

Our current hypothesis on the role of charged regions in LLPS is that these can provide structural/dynamical features that can be robustly maintained both in the solution and the condensed phase and help the relative positioning of the regions responsible for specific interactions. The possible more direct role of the charged motifs in actual condensates can also not be excluded but would require complex experimental investigations.

## Methods

### Surveying protein sequences for charged residue repeats

The version of the human reference proteome from UniProtKB (https://www.uniprot.org/proteomes/UP000005640) used for this study encompasses 20 659 human genes - with one isoform per gene - and represents the 2020 September 3 update. Some investigations involved the exclusion of transmembrane proteins that was carried out by removing entries from the above mentioned fasta file based on another UniProtKB query that encompassed 41 818 sequences - that aren’t necessarily part of the reference proteome - annotated as human transmembrane proteins (the exact query was annotation:(type:transmem) AND organism:“Homo sapiens (Human) [9606]”). Furthermore, additional small sequence sets matching the length distribution in PhasepDB have been generated by randomly selecting one - or ten - from the above mentioned 20 659 sequences for each (human) PhaSepDB entry where the two must have had a similar (±5%) length.

To identify segments with regular charge patterns (CRRs), the method FT_CHARGE was used. FT_CHARGE was developed to identify SAH regions in proteins that contain a repeating charge pattern with a specific frequency between 1/9 and 1/6 but it is also capable of detecting other motifs with regularly alternating positively and negatively charged residues. [47] In this work the largest window size for FT_CHARGE was 64 residues, corresponding to a minimum frequency of 1/64. The frequency of 1/64 practically corresponds to a long, repeated segment of identically charged amino acids (e.g., a polylysine run), whereas the frequency of 1/2 corresponds to a region of residues with alternating charges (e.g., the sequence “KEKEKEKEKE”). Such detected sequences are hereafter denoted as “charged residue repeats” - or CRRs for short, to make a distinction between these and “genuine” SAHs. In this view, SAHs represent a subset of CRRs. For more details on the implementation of FT_CHARGE refer to [48, 49].

Detailed data for each sequence and statistics on CRRs are shown in Supplementary Tables S7, S7 and on Supplementary Figure S10.

### Clustering human proteins based on structural predictions

To remove redundancy from the sequence set, the webserver CD-HIT was employed to cluster the sequences on their similarity. The cut-off values for this were 0.9, 0.7 and 0.5, meaning 90%, 70% and 50% sequence identity, respectively. Clustering was carried out with global sequence identity, a 20-residue bandwidth of alignment, a minimal sequence length of 10 residues, and default alignment coverage parameters. Sequences were assigned to the best cluster that met the threshold. The output of CD-hit was further processed using MATLAB to ensure compatibility with our analysis pipeline. The three redundancy-filtered arrays obtained have contained clusters of UniProt IDs with a pre-set cut-off value (90%, 70% and 50% sequence identity).

### Surveying protein sequences for Charge-Dense Regions (CDRs)

While SAHs - and CRRs in general - may contain a higher-than-average amount of charged residues, their features mainly result from the specific patterns these residues repeat within the sequence. Thus, a separate set of investigations must be carried out if we are to expand the analysis of the correlation between charged protein regions and phase separation. We applied two alternative approaches for identification of such regions, the first one considers a region highly charged if it contains a higher-than-average ratio of charged residues, while the other identifies sub-sequences whose overall charge significantly differs from zero. Based on this difference the recognized “Charge-Dense Regions” (CDRs) will be called “signed” and “unsigned” CDRs, or sCDR and uCDR motifs, respectively.

The scoring scheme for sCDR assigns a score of +1 to arginines, histidines and lysines, and −1 to aspartic and glutamic acids, while the one for uCDRs assigns a score of 1 to all five of the aforementioned residues. In both cases, the investigated sequence window is considered to contain a CDR if the absolute value of its overall score divided by its length reaches a pre-set cut-off value.

To determine adequate cut-off values for all the used window sizes, we run the algorithm on a randomized version of the human proteome with a cut-off value of zero. The randomized sequences had the same amino acid composition and length distribution as the natural ones. Based on the obtained scores, two cut-off values were determined, one that yields roughly 5% of all hits and one that yields only about 1% - note that the discrete nature of scoring (i.e., steps of 1/16 between 0 and 1 for a window length of 16 residues), selection of a threshold is somewhat approximated. Using these two cut-offs, the full proteome was scanned to identify signed and unsigned CDRs in the wild-type sequences.

The segments identified using different sequence windows were merged for each sequence. During the process, care was taken to ensure that the merged segments are still above the predefined threshold, which needs special treatment for signed CDRS where the overall charge should also be maintained. In the final step, all data were unified for all window sizes used, 16, 32, 64 and 128.

### Conducting Fisher’s exact test of independence on members - and clusters - of the human proteome

To assess the association of specific sequence sets with LLPS, 2×2 contingency tables were generated and then Fisher’s exact test as implemented in R [R statistics package reference] was applied to determine the P-value.

In case of clusters, two different methods were applied for categorization. In the first one, evaluation is applied cluster-wise, meaning that association with LLPS for a given cluster was considered positive if any member (sequence) in the cluster was annotated as such in PhaSepDB. In the second, a cluster was only considered to be participating in LLPS if the representative sequence kept by CD-HIT was annotated in PhaSepDB. The presence or absence of CRRs was established in an analogous manner.

### Preparing receiver operating characteristic (ROC) tests

The purpose of ROC tests is to evaluate an attribute as a viable indicator for classification. In the case of this study, they were used to investigate the presence of different sequence motifs as indicators for a protein’s affinity towards liquid-liquid phase separation. We have sorted the proteins in the human proteome according to their score obtained in the CDR/CRR detection and calculated the true positive, true negative, false positive and false negative rates.

## Author Contributions Statement

Z.G. and P.R. designed the study, A.L.S. developed software, A.L.S and A.S. analysed the data, all authors contributed to data interpretation, drafting the article and approved the final version.

## Acknowledgements

This work was supported by a grant from the National Research, Development and Innovation Office under grant OTKA 124363 to Z.G. and FK-128133 to R.P., and a fellowship (grant number 158534) from the Tempus Public Foundation of the Hungarian government to R.P. A.S. acknowledges the support by an ÚNKP Fellowship.

