## Supplementary tables and figures for "Charged sequence motifs increase affinity towards liquid-liquid phase separation": Szabo_ChargedMotifsLLPS_SupFigs.pdf

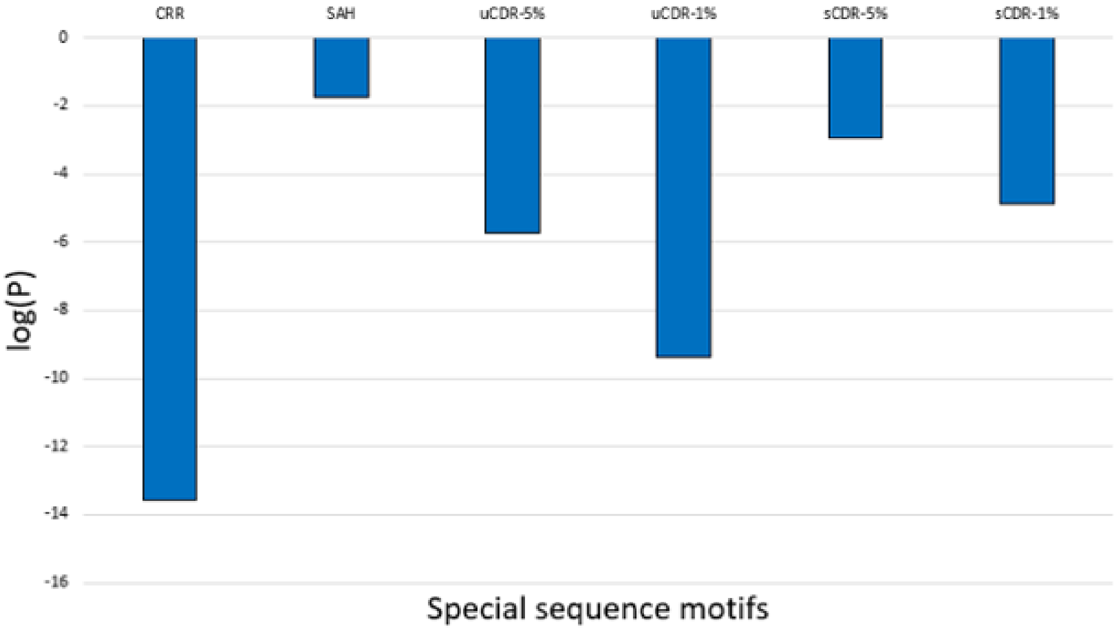


**Figure S1.** Decimal logarithm of P-values obtained from Fisher's exact test of independence, carried out respectively on pairs of attributes where one attribute was the association to LLPS (whether the sequence was present in PhaSepDB) and the other attribute was the presence of a given type of charged sequence motif. In case of CDRs these tests were conducted with two thresholds for the motifs, in one case the motifs had to be within the top 1% of hits while in the other case it was enough if they were in the top 5% of hits.

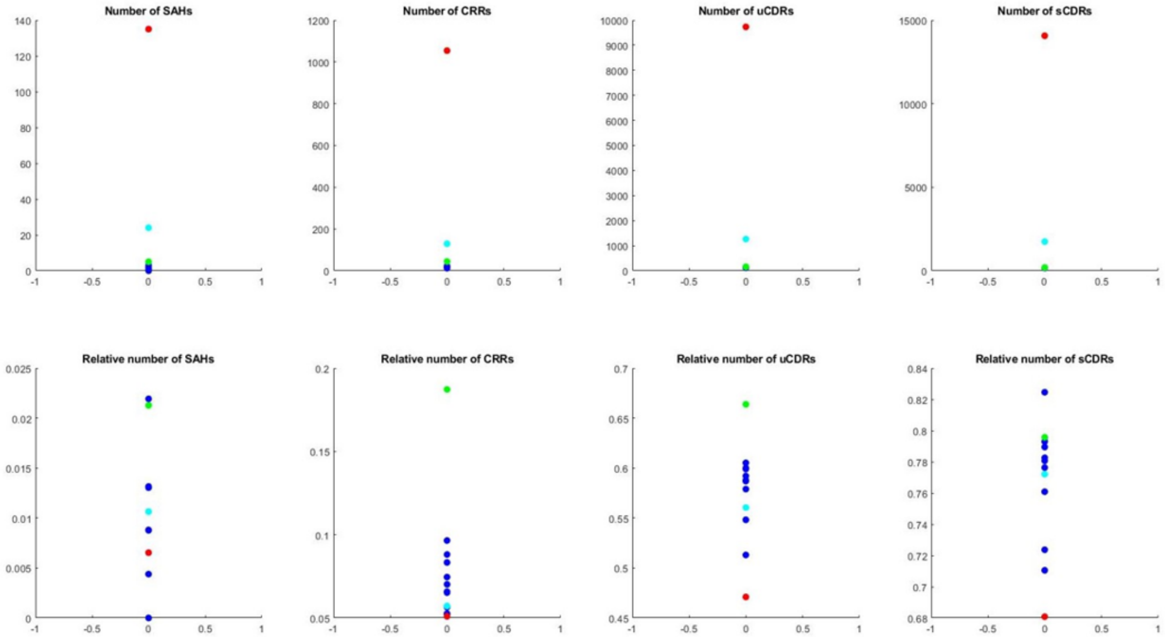


**Figure S2.** Number (top row) and fraction (bottom row) of sequences with charged motifs within the human reference proteome (red), PhaSepDB (green), ten smaller sequence sets where one sequence with a similar length (5%) has been randomly selected for each human PhaSepDB entry (blue), and one sequence set with ten randomly selected, similarly long sequences per PhaSepDB entry (light blue). In the random selection process, already selected sequences were removed from the pool of possible choices, for the sake of minimizing redundancy.

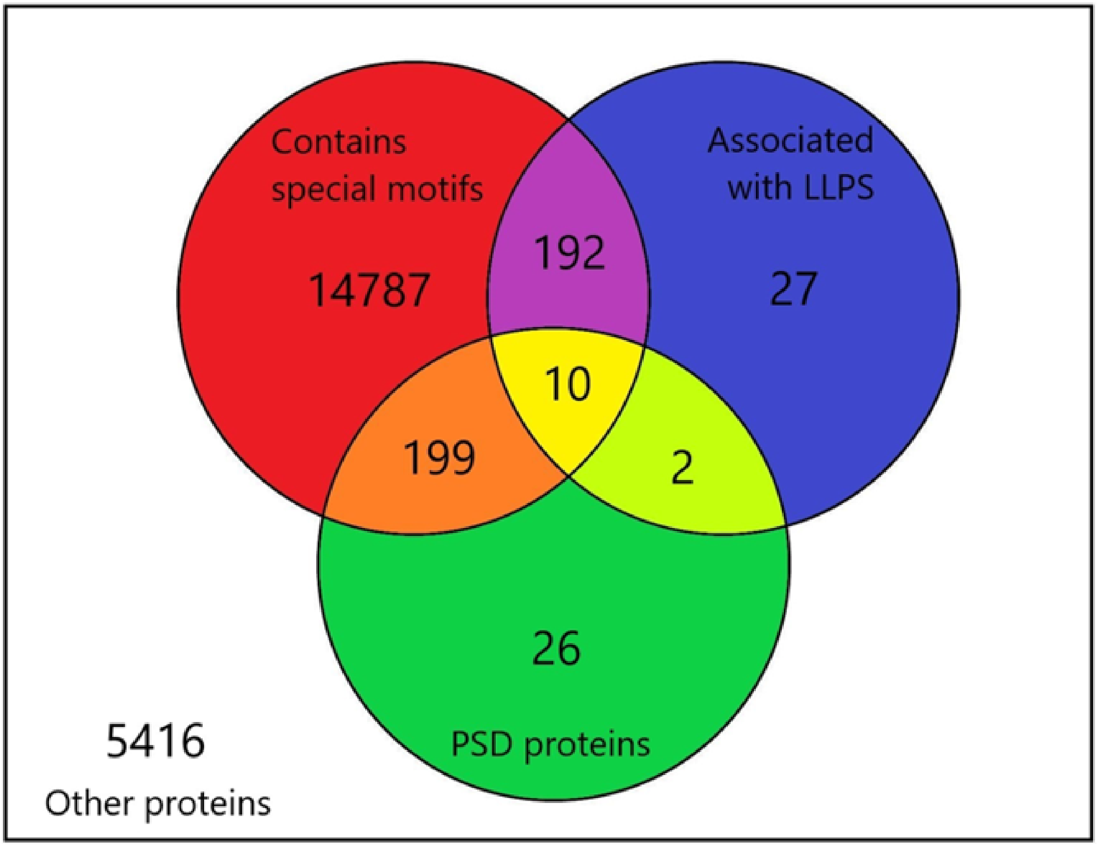


**Figure S3**. Categorization of sequences based on their charged motif-content, association to LLPS and relation to the postsynaptic density (PSD).

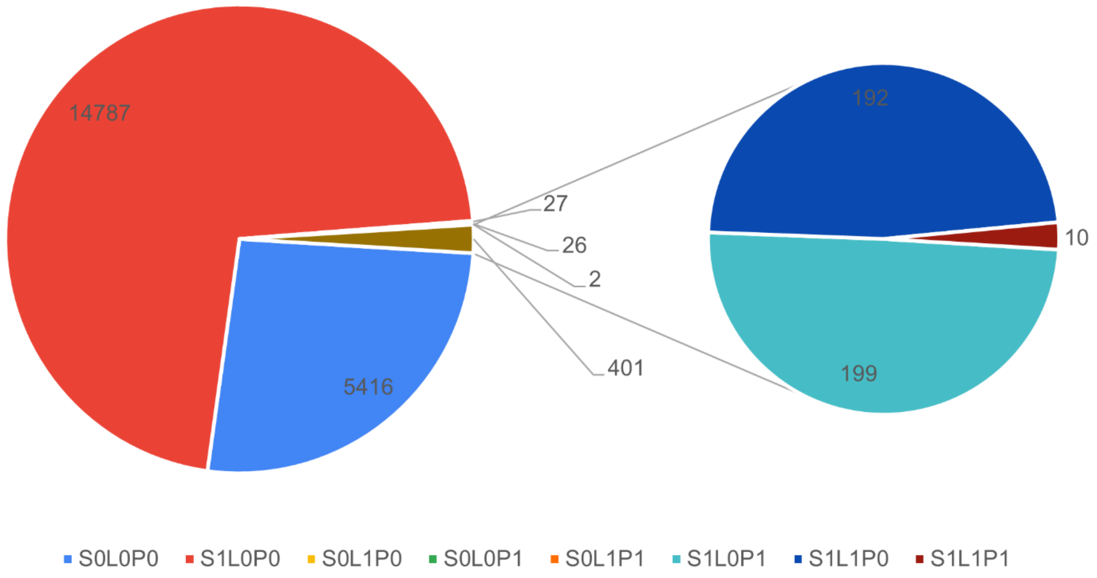


**Figure S4**. Number of entries within each category shown on Figure S3. Entries were categorized based on whether they contain any charged sequence motifs (S0/S1), whether they are associated with LLPS (L0/L1) and whether they are related to the postsynaptic density (P0/P1).

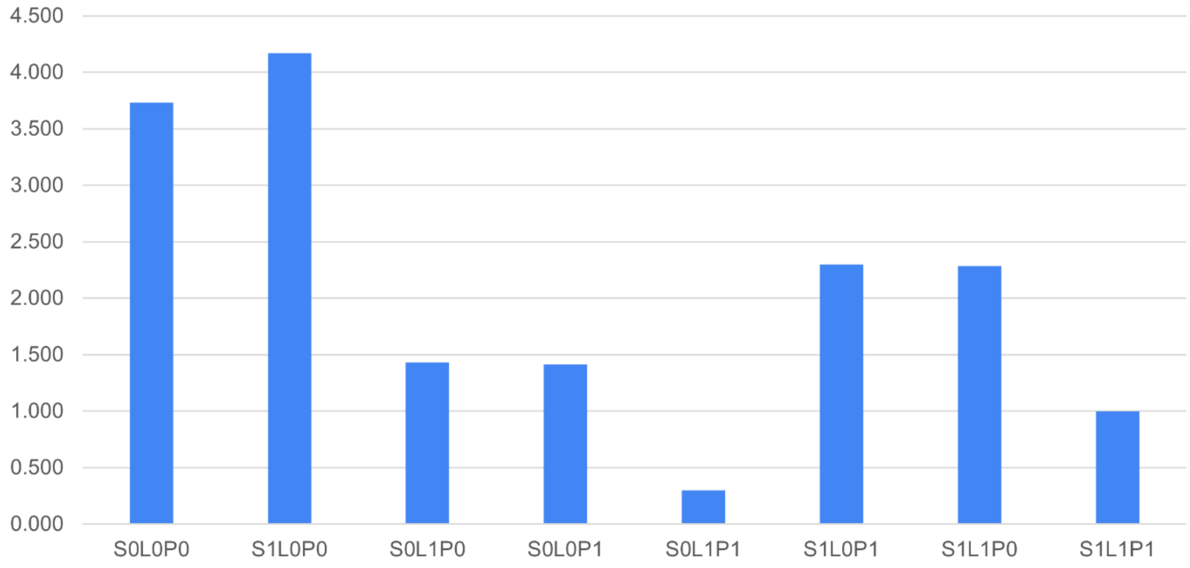


**Figure S5.** Decimal logarithm of the number of entries within each category shown on Figure S3. Entries were categorized based on whether they contain any charged sequence motifs (S0/S1), whether they are associated with LLPS (L0/L1) and whether they are related to the postsynaptic density (P0/P1).

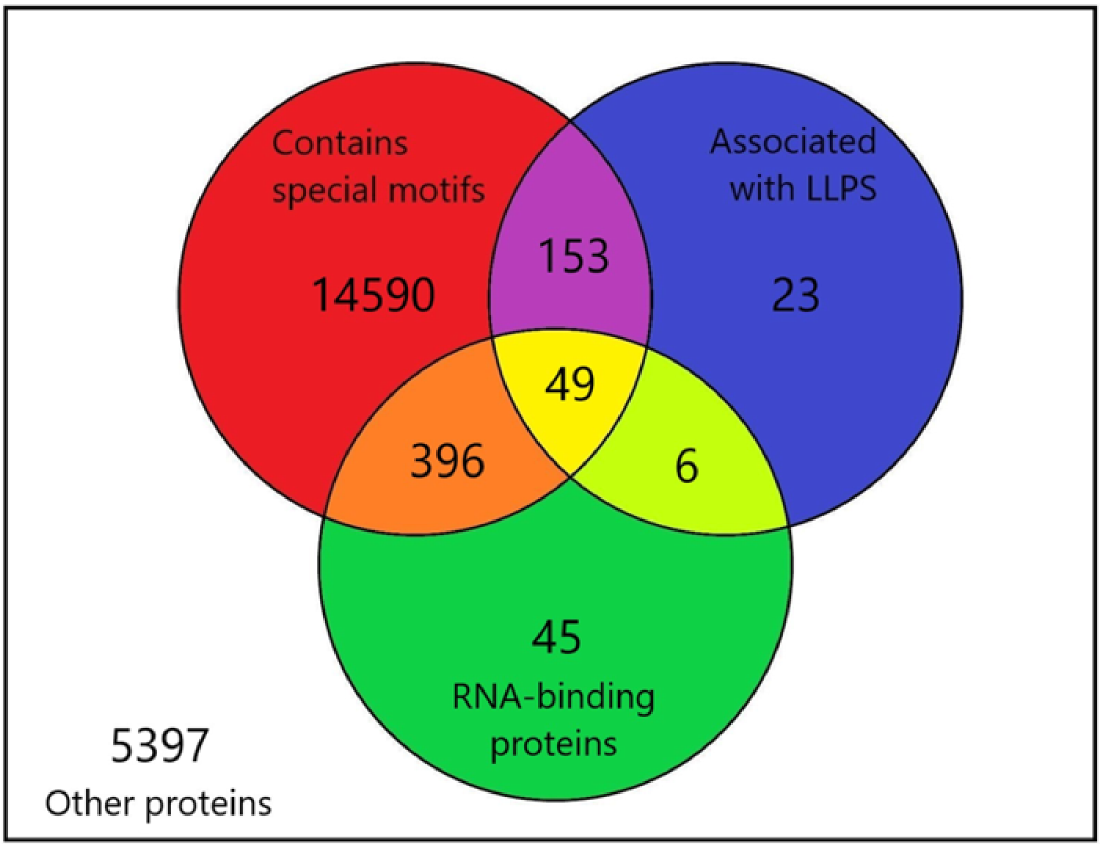


**Figure S6.** Categorization of sequences based on their charged motif-content, association to LLPS and ability to bind RNAs.

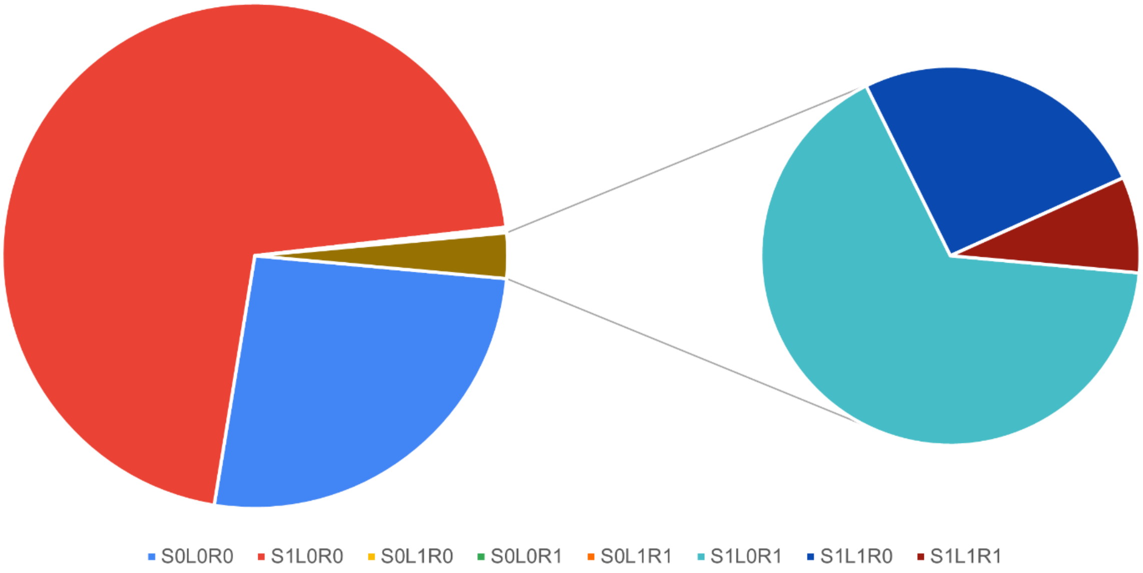


**Figure S7.** Number of entries within each category of Figure S6. Entries were categorized based on whether they contain any charged sequence motifs (S0/S1), whether they are associated with LLPS (L0/L1) and whether they are known to bind RNAs (R0/R1).

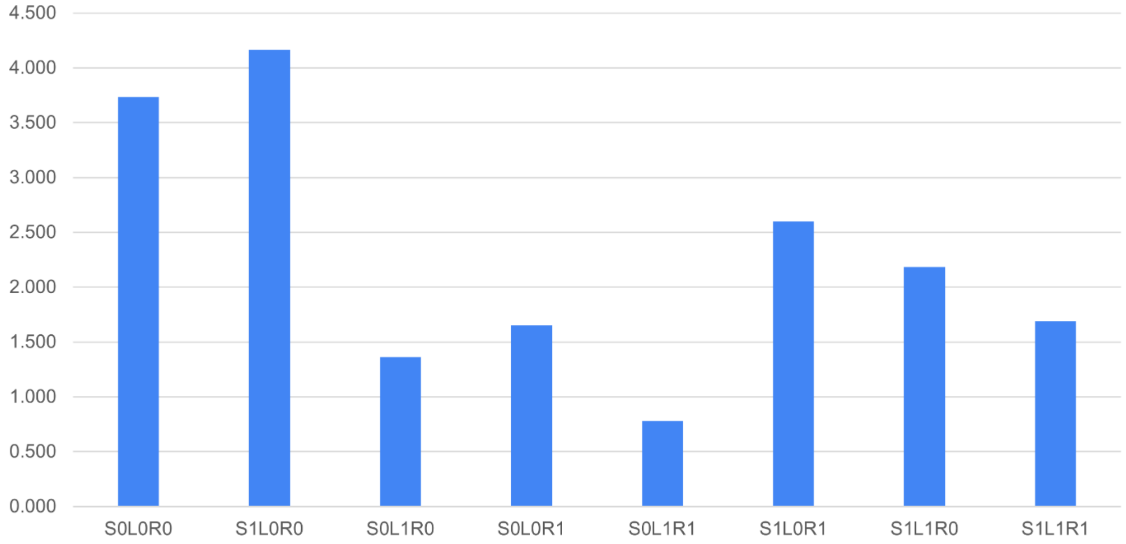


**Figure S8.** Decimal logarithm of the number of entries within each category of Figure S6. Entries were categorized based on whether they contain any charged sequence motifs (S0/S1), whether they are associated with LLPS (L0/L1) and whether they are known to bind RNAs (R0/R1).

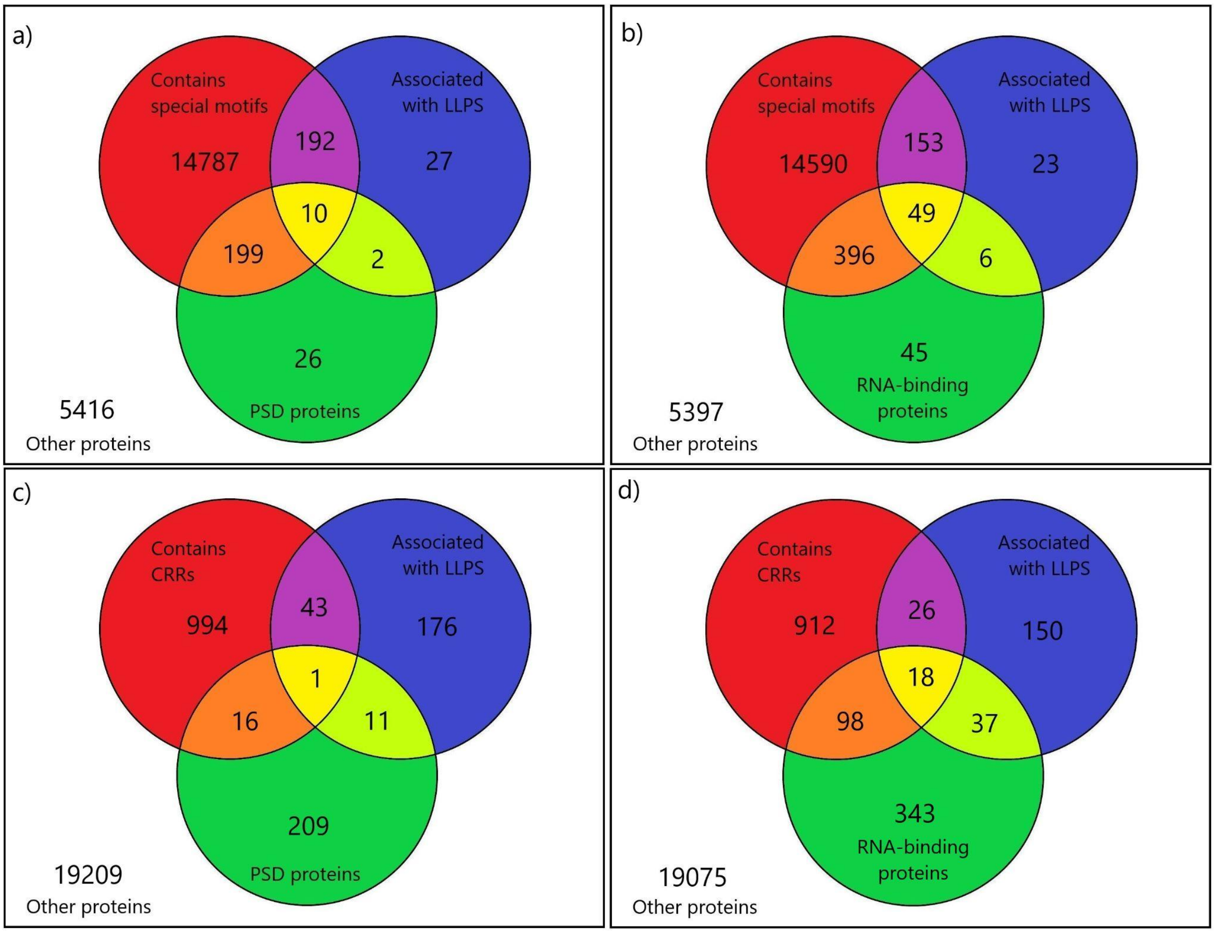


**Figure S9.** Categorization of all human proteins (including transmembranbe proteins) according to their special charged motif- (a-b) or CRR-content (c-d), LLPS-association and their functional role in the PSD (a, c) or RNA-binding (b, d).

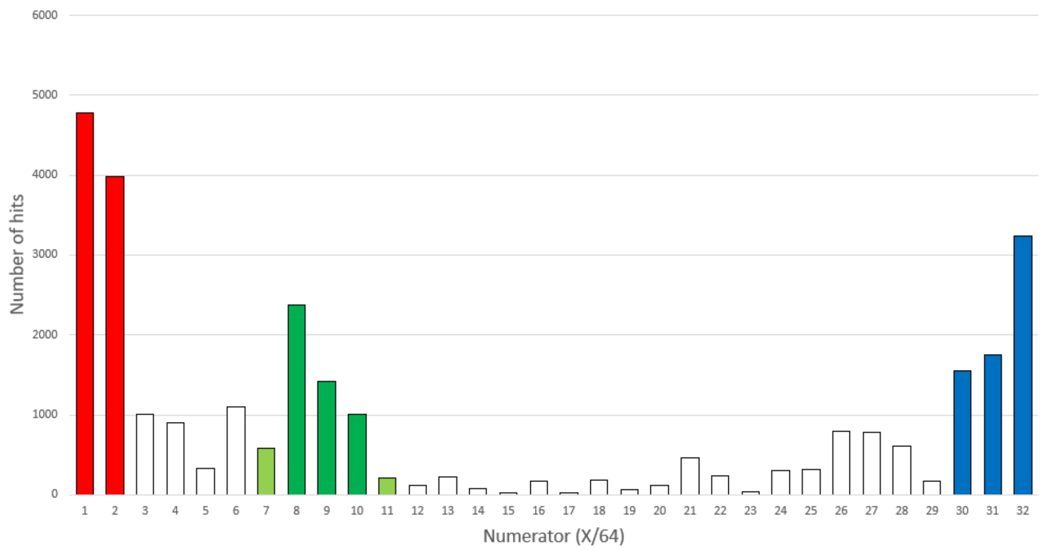


**Figure S10.** Number of FT_CHARGE hits organized by their Fourier frequency (1/64 - 32/64). Three separate groups can be distinguished by their prominent number of hits: 1/64-2/64, where charged residues are quite scarce, 30/64-32/64, where about every other residue is charged, and 8/64-10/64, which is within the range of SAHs (1/9-1/6). Hits with frequencies of 7/64 and 11/64 are just outside this range and are therefore omitted from the group.
